# Somatic evolution of cancer genes on the sex chromosomes

**DOI:** 10.1101/2024.09.11.612525

**Authors:** Amarachi C. Akachukwu, Jasmine S. Ratcliff, Greg N. Brooke, Antonio Marco

**Affiliations:** School of Life Sciences, University of Essex, Wivenhoe Park, Colchester CO43SQ, United Kingdom; Institute for Social and Economic Research, University of Essex, Wivenhoe Park, Colchester CO43SQ, United Kingdom

**Author notes:** Corresponding author (AM). These authors contributed equally to the work (Amarachi C. Akachukwu, Jasmine S. Ratcliff).

## Abstract

Cancer progression arises through the somatic evolution of oncogenes (OGs) and tumour-suppressor genes (TSGs). According to Knudson’s two-hit hypothesis, both alleles of a TSG must typically be inactivated for tumorigenesis to proceed, whereas a single activating mutation can drive an OG. However, this diploid framework does not apply to genes in the male X chromosome, which is hemizygous, so that a single loss-of-function mutation can inactivate an X-linked TSG. Here, we adapt classical population genetics models of adaptation in haploid and diploid asexual populations to somatic evolution, and we test their predictions using somatic mutations from the COSMIC database. After excluding mutations in genes with reported copy-number alterations or promoter methylation, we find that TSGs in tumours from male patients show a higher ratio of loss-of-function (nonsense and frameshift) to synonymous mutations than autosomal TSGs, and that complete losses of X-linked TSGs are more frequent in males. Importantly, we find no such differences in tumours arising in females. X-linked OGs also show a higher rate than autosomal OGs in males and, as predicted by the model, the pattern is reversed in low-proliferation tumours, although this last difference is not statistically significant. Neutral (synonymous) mutation rates differ between sexes but not between the X chromosome and autosomes. Our work shows that ploidy and chromosomal context can be incorporated into models of somatic evolution, and it provides a framework to explore sex differences in tumorigenesis.

## INTRODUCTION

Cancer results from the abnormal growth of populations of cells that escape multiple cellular control mechanisms [1,2]. Cancer development is a multistage process and typically requires one or more genes to be mutated at the somatic level [3–5]. A somatic mutation in a cancer-associated gene that is detrimental to the host may confer a selective advantage on tumour cells by promoting their proliferation, survival or dissemination. Traditionally, two major types of cancer genes have been identified, depending on the type of mutation required to benefit the progressing tumour: oncogenes and tumour-suppressor genes. Oncogenes (OGs), or proto-oncogenes when they are normally functioning in the host, are usually activated by gain-of-function mutations, typically dominant, as only one of the two copies needs to be mutated to have an impact. On the other hand, tumour-suppressor genes (TSGs) usually require both copies to be inactive (by e.g. mutation, deletion, or silencing) to have an effect on cancer, and therefore these mutations follow a traditional recessive inheritance pattern. The Knudson two-hit hypothesis states that a TSG involved in cancer development should have both of its alleles inactivated, either by mutation or by another mechanism [6]. We hypothesize that the chromosomal context of cancer genes (sex or autosomal) and the type of activating mutation (gain or loss of function) significantly influences how fast damaging mutations leading to cancer accumulate in the genome.

Sex differences in cancer incidence, outcome and molecular profiles are well documented [7–12]. Males show a higher cancer risk and an earlier onset in many tumour types, and sex differences have been described in oncogenic mutational processes [13] and in the landscape of somatic alterations [14] . Part of these differences has been attributed to genes on the X chromosome. In males, X-linked genes are hemizygous, so a single loss-of-function mutation can inactivate an X-linked TSG, whereas in females most X-linked TSGs are expected to need two hits, with the exception of those that escape X-inactivation [15]. This has also been proposed as an explanation for the evolutionary depletion of TSGs on the X chromosome [16]. Hemizygosity, however, is not equivalent to loss of heterozygosity (LOH) in autosomal TSGs. LOH, which can arise through deletion, mitotic recombination or chromosome loss [17,18],represents an additional event following the initial mutation, with its own probability of occurrence and timing. In contrast, a single loss-of-function mutation in an X-linked TSG in males may be sufficient to abolish gene function. These studies have mostly examined how often TSGs are altered. Here, we take a complementary approach: we use population genetics models of adaptation in haploid and diploid asexual populations to predict the relative rate of adaptive mutations in TSGs and OGs, in the X chromosome and autosomes, and as a function of the size of the tumour cell population.

To understand the dynamics of genetic mutations in cancer development, multiple mathematical models have been developed during the last decades. Cancer somatic evolutionary models are often based on the Luria-Delbrück model or its derivations (reviewed in [19]). These models often implement realistic parameters to account for cancer complexity, such as explaining the different rates of cancer spread under different conditions. For instance, recent dynamic models implement multistage processes [20,21], compartmentalization [22], the joint impact of pro- and anti-cancer mutations [23], or even epistatic interactions [12]; see [25] for a historical overview of the development of cancer evolution models. These models of cancer evolution often assume full diploidy of the cells (or just ignore ploidy), which may have an impact when studying sex chromosomes. Instead, here we propose the application of traditional population genetics models that account for chromosomal ploidy and dominance of mutations in somatic evolution.

The rate at which beneficial mutations are fixed in a population has been a central topic in population genetics. If beneficial mutations are fixed in one gene more often than in another gene, we say that the first gene evolves faster than the second. One classic result is that, overall, genes in haploid organisms evolve comparatively faster than those in diploid organisms [26,27]. This is because the majority of newly introduced mutations are recessive with respect to the wild-type allele in diploid populations, and the beneficial effects of the new mutant are not manifested in heterozygosis. This result can be extended to the relative evolutionary rate of genes in autosomes and sex chromosomes. A gene located on an X chromosome in mammals, when in males, is effectively in haploidy (hemizygosis). Theoretical models have shown that, in the long term, genes located on the X chromosome should evolve faster than genes located on autosomes: the faster-X effect [28]. However, some contradictory data related to the faster-X effect exists [29], as well as a number of confounding factors [30] that should be taken into account. In any case, these models assume diploidy and sexual reproduction so, in the context of somatic evolution in cancer, asexual models for both haploids and diploids should be considered in order to understand a potential impact of the chromosomal context.

Some early models comparing haploid and diploid systems were developed by Crow and Kimura [31] based on earlier work in theoretical population genetics [32,33]. These models compared the evolution of genes in haploid and diploid (sexual) systems, concluding that sexual reproduction results in faster gene evolution. They also proposed that diploidy may confer some protection against damaging mutations in somatic tissues. This theory was further progressed by Orr and Otto [34] who, building upon the Crow-Kimura-65 model, specifically compared the rate of emergence of adaptive mutation between haploids and diploids in non-recombining systems. They conclude that, in somatic (asexual) evolution, the rate of adaptation depends, not only on the rate of emergence of new favorable mutations (ploidy) but also on the rate at which these mutations are fixed (dominance and population size).

We aim to investigate whether there is a difference in the rate of adaptive mutations in either TSGs and/or OGs between sex chromosomes and autosomes and the relative role of selective pressure and somatic mutation rates in their evolutionary rates during cancer progression. To do so, we build upon Crow-Kimura-65 and Orr-Otto-94 models, considering ploidy as a proxy for chromosome location, and evaluating our predictions by analyzing whole genome mutation screens from multiple cancer samples.

## RESULTS

### The rate of adaptation of cancer genes in a chromosomal context

To evaluate the rate of adaptation we consider the rate of emergence of a favorable mutation escaping stochastic loss per individual and per generation, *U*. Crow and Kimura [31] showed that, in asexual organisms, the average time to fixation of a favorable mutation is *1/U* and given by:

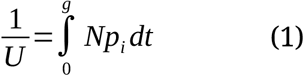

where *p_i_* is the frequency of the advantageous mutation at time *t*, *g* is the average number of generations that it takes to reach *p_t_*, and *N* is the effective population size. By assuming that the frequency of the advantageous mutation follows a logistic growth [35], they found that:

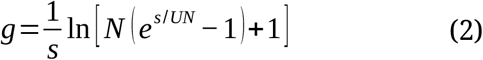

where *s* is the selective advantage of the mutant for asexual haploid populations, yet this term is *hs* for diploid asexual populations [34] where *h* is the dominance coefficient of the mutant with respect to the wild-type allele.

Orr and Otto [34] build upon this model and approximate forms of *g* for haploid and diploids separately and, since the rate of adaptive evolution (*K*) is inversely proportional to the average time to fixation (*g*) they derived that:

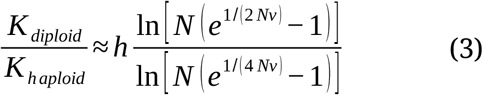

where *v* is the “rate at which favorable alleles appear by mutation per haploid genome” [34]. From here we will refer to the ratio K_dip_/K_hap_ as *R*, the relative rate of adaptive evolution of diploids with respect to haploids. Hence, a value of *R* greater than 1 indicates a faster evolution in a diploid context, and a value less than 1 indicates a faster evolution in a haploid context.

In the context of cancer somatic evolution, cancer genes on the X chromosome in male tissues are modeled as haploids, whilst autosomal genes (and X genes in female tissues, with some caveats that we will explore later) are modeled as diploids. Further, oncogenes (OGs) are characterized as becoming activated by gain-of-function dominant mutations (we explore this below), and hence we can assume partial or full dominance: *h* > 0.5. On the other hand, and following Knudson’s two-hit model [36], Tumour Suppressor Genes (TSGs) are characterized for loss-of-function recessive mutations, so we assume either full or partial recessiveness: *h* < 0.5. Orr and Otto gave approximate solutions for the case of rare favorable mutations (*R*≈*2h* for *Nv* << 1) and for more frequent favorable mutations in the population (*R*≈*h* for *Nv ≥* 1).

### Estimation of the parameter *Nv* in human tissues

It is challenging to estimate *Nv* in human somatic tissues and tumours, yet we can attempt a rough approximation. The estimation of somatic mutation rates varies from study to study, yet most agree that it is in the range of 10^-8^ to 10^-9^ mutations per nucleotide per cell division [37–39]. Assuming that ∼1.5% of the nucleotides of the human genome are part of codons, we can set up an upper limit (as not all mutations in codons are non-synonymous and beneficial for cancer progression) at 0.015 as the proportion of mutations that are beneficial to tumour growth. Hence, *v* may be in the range of 1.5 x 10^-10^ to 1.5 x 10^-11^ beneficial substitutions per nucleotide per cell division.

The estimation of *N* is also problematic as different tissues have different sizes, and also only a fraction of a tissue is usually proliferative. We can start by considering skin, the largest organ that has a high proliferative rate. It is estimated that there are approximately 2.7 million proliferative cells per cm^2^ of skin under normal conditions [40]. Given that the average size of the skin surface in an adult is around 1.6 m^2^ [41], that gives us a total of 43.2 billion of proliferative cells. Given these estimates, the *Nv* in skin should be between 0.65 and 6.5. We can consider now a smaller organ with a lower proliferative rate, like in the case of the pancreas. Estimates from healthy individuals report an average number of acinar cells of 1.22 x 10^11^ cells per adult pancreas [42]; acinar cells are largely responsive for the development of carcinomas [43]. Although less than 2% of these cells are proliferative at the same time in an adult pancreas [44,45], this still gives an estimate of *Nv* between 0.037 and 0.37.

However, these rough estimates are more indicative of the rate of emergence of pioneer/driver mutations in healthy tissues. Once a tumour clone is established, the value of *Nv* during cancer progression will depend on the effective population size of the tumour and its mutation rate, which can be estimated using specialised computational tools [46]. Consequently, tumours with lower proliferative activity may exhibit lower Nv values. In this context, N refers to the effective population size (N_e_), rather than the total number of tumour cells. For instance, in Clear Cell Renal Cell Carcinoma (ccRCC) is has been estimated that there are approximately 1.13 x 10^5^ cells per mm^3^ of tumour tissue [47]. For a developing clonal tumour population with a volume of 1 cm3, *Nv* is estimated to range from 0.0017 to 0.0170, assuming no acceleration in the mutation rate. According to the model described above, this suggests that genes located on the sex chromosomes may exhibit different evolutionary dynamics in tumours with low proliferative activity compared with those exhibiting high proliferative activity. We explore these differences further in this study.

If the model discussed holds for somatic cancer evolution, we can make the following precise predictions:

(1) In male mid- and high-proliferative tumours (*Nu ≥* 1), advantageous mutations accumulate (collectively) faster in the X chromosome than in autosomes, with this effect being stronger in TSGs (h<1/2).
(2) In male low proliferative tumours (*Nu* << 1), advantageous mutations in TSGs accumulate faster in the X chromosome than in the autosomes (h<1/2) and vice versa in OGs (h>1/2).

### Time to fixation

In the above model, the Poisson process with rate *U*, the time to advantageous mutation occurrence is exponentially distributed [48] and, therefore, the expected (average) arrival time is *1/U*. This is precisely where the term *1/U* in equation (1) comes from [31]. Therefore, the expected time for an advantageous mutation to occur in haploids is:

and for diploids it is:

According to the equation above, the time to advantageous mutation arrival is dependent on the coefficient of dominance in diploids, where for dominant mutations (h>0.5) the time to arrival is smaller than for recessive mutations (h<0.5). In this context, ‘fixation time’ refers to a parameter from the evolutionary model that doesn’t have a straightforward meaning in the context of somatic evolution, but that it is here interpreted as interpreted as a proxy for the time to clonal dominance. From the equations here described we can make a third prediction from the model, in addition to the two made in the previous subsection:

3) Advantageous mutations arise, on average, earlier in OGs than in TSGs, except for X chromosomes in hemizygosis, where both arrise, on average, at the same time.

### The nature of mutations in cancer genes

The classification of cancer genes into OG or TSG is a simplification. However, it often captures the dominance of the most likely driver mutations. From the COSMIC dataset here used we identified the number of OG and TSGs that are classified as having recessive or dominant mutations in cancer screening (Table 1, Supplementary Table 1). We also include genes classified as both TSG and OG and also those reported to be both dominant and recessive. From the data available it is evident that oncogenes are mostly characterized by hosting dominant (probably gain-of-function) somatic mutations, while TSGs are more likely to host recessive mutations (p < 0.001; Fisher’s Exact test).

**Table 1.** Reported somatic mutations in census cancer genes from the COSMIC database.

|  | Dominance |  |  |
| --- | --- | --- | --- |
|  | Dominant | Recessive | Both |
| **OG** | 204 | 3 | 0 |
| **TSG** | 71 | 117 | 2 |
| **POG and TSG** | 40 | 16 | 5 |

It is expected that dominant mutations are mostly gain-of-function and recessive are loss-of-function. Looking at coding genes, gain-of-function mutations like those expected in OGs should mostly involve a change of an amino acid (although this is not always the case, see Discussion). On the other hand, we will expect an enrichment of nonsense mutations (gain of a STOP codon) as loss-of-function somatic mutations and, therefore, in TSGs. We compare the number of nonsense and missense (nonsynonymus) mutations identified in genomic screens (see Methods). As expected, the number of nonsynonymous mutations is proportionally higher that the number of nonsense mutations in OGs (Figure 1; slope = 0.75), and the opposite is true for TSGs (Figure 1; slope = 1.19). Both regression lines are statistically independent (p < 0.001; regression interaction). For the study of TSGs, despite losing some statistical power, it is convenient to consider only nonsense mutations to avoid including gain-of-function nonsynonymous mutations that may bias the outcome. We conclude then that, as a good approximation, TSGs are characterised by loss-of-function recessive nonsense mutations whilst OGs are characterised by gain-of-function dominant nonsynonymous mutations.

**Fig 1.**
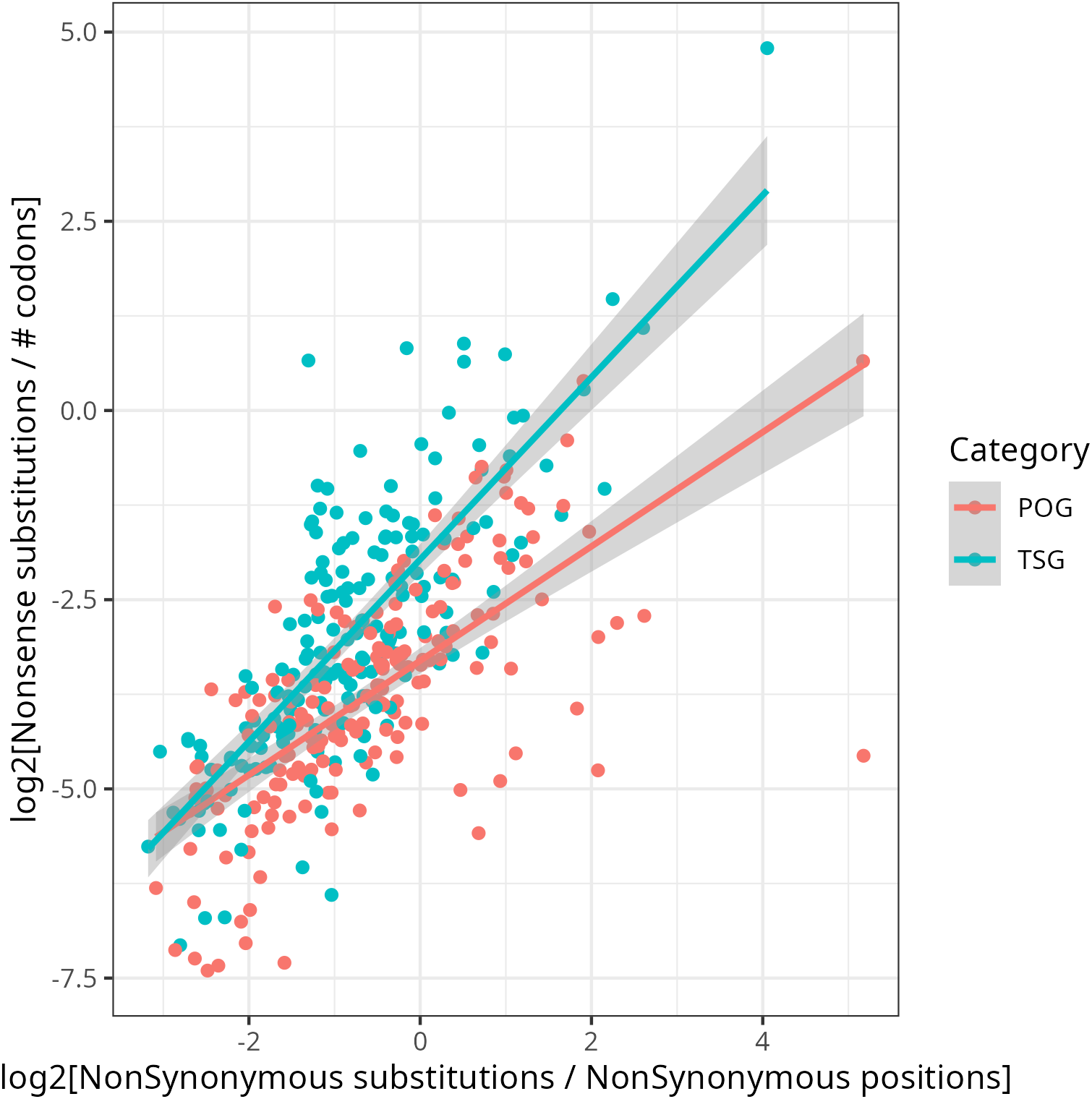
between nonsynonymous and nonsense somatic mutations in cancer genes. between nonsynonymous and nonsense somatic mutations in cancer genes. Scatter-plot of the normalised number of non-synonymous somatic mutations versus the number of nonsense (STOP codon gain) somatic mutations mutation in the Genome screens in the COSMIC database. Data shown for proto-oncogenes (POG) in red and for tumour-suppressor genes (TSG) in red. Lines of the same colour are the linear model fit, including the error for each line.

### Evolutionary rates at the sex chromosomes

As a proxy for the somatic evolutionary rate of cancer genes we compute the log-odds ratio (LOR) as the logarithm (natural) of the ratio of the fraction of function changing mutations over potential function changing sites with respect to the fraction of synonymous mutations over synonymous sites (see Methods for details). For TSGs we considered STOP codon gain mutations are change of function, while for OG we consider nonsynonymous mutations (see rationale in the previous section). The genomic screen studies were classified according to whether the cancer samples were retrieved from females or male samples, and the analysis were performed separately for both sexes.

Importantly, copy number variations are frequent somatic changes associated with cancer[49]. This will alter the local ploidy and, therefore, potentially affect our analysis of single-nucleotide somatic mutations. To avoid this, we excluded mutations in genes for which copy number alterations (either gains or losses) had also been reported in the same study. About 5% of the mutations in the genomic screens were filtered out. Likewise, we also remove those genes in samples with reported differential methylation in their promoters (see Methods), although those only accounted for about 0.3% of the total dataset.

In TSGs, the LOR (proxy for evolutionary rate) in female cancer samples was comparable between autosomes and the X chromosome (Fig 2A, p = 0.238; Mann-Whitney test). However, in male samples, and in concordance with the theoretical model described above, the evolutionary rate was significantly higher in genes in the X chromosome compared to genes in autosomes (Fig 2B, p = 0.022). The effect size of this difference is notable (difference between the medians = 1.04). In addition, we considered frameshift mutations as another type of mutation causing loss-of-function. Consistently with the results observed for nonsense mutations, the evolutionary rate based on frameshift loss-of-function mutations was not distinguishable between X and autosomes in female cancer samples (Figure 2C), yet it was statistically higher in the X chromosome compared to the autosomes in male samples (Figure 2D).

**Fig 2.**
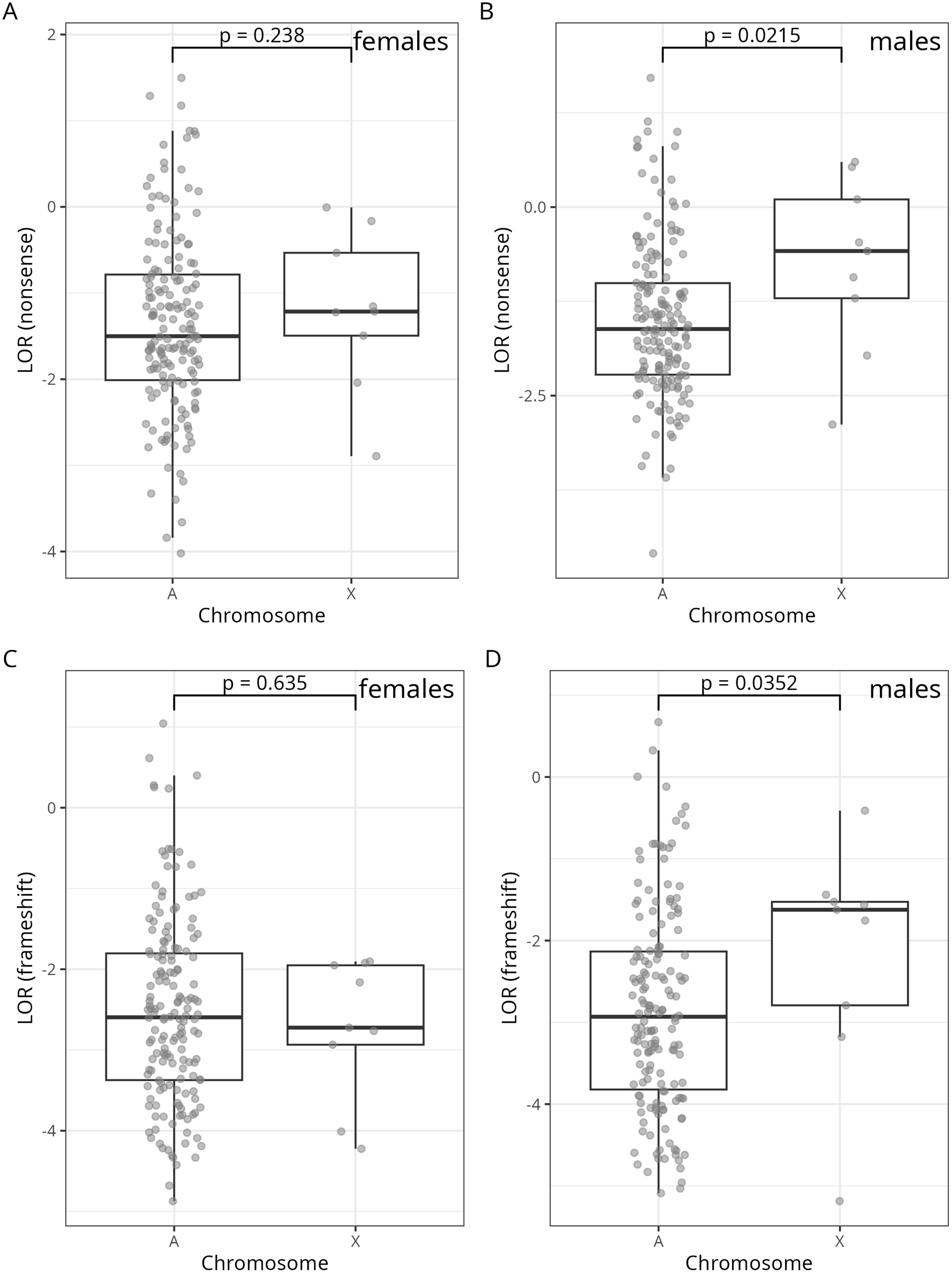
Loss-of-function mutations in TSGs for males and females in autosomes and the X chromosome. Boxplot of the scaled mutation rate (log-odds ratio (LOR) of the number of loss-of-function mutations, see Methods). The top panels plot the LOR based on nonsense mutations of TSGs located in any autosome (A in the x-axis) or in the X chromosome (X in the x-axis) for female (A) and for male (B) cancer samples. The bottom panels plot the LOR based on frameshift mutations of TSGs located in any autosome or in the X chromosome for female (C) and for male (D) cancer samples.

In OGs, like with TSGs, the evolutionary rate of genes in autosomes is comparable to that of genes in the X chromosome in female cancer samples (Fig 3A, p = 0.585). In male samples, and again in agreement with the theoretical model, genes in the X chromosome evolve faster compared to genes in the autosomes (Fig 3B, p = 0.018). A specific prediction of the model is that the relative accelerated evolution of genes in the X chromosome with respect to autosomes should be stronger in TSG compared to OG. Indeed, the effect size of the differences in evolutionary rate in OGs is smaller, as predicted (difference of the medians = 0.3912) than in the case of TSGs. We conclude that the empirical data available from the study of cancer samples agrees with the first prediction we made from the theoretical model.

**Fig 3.**
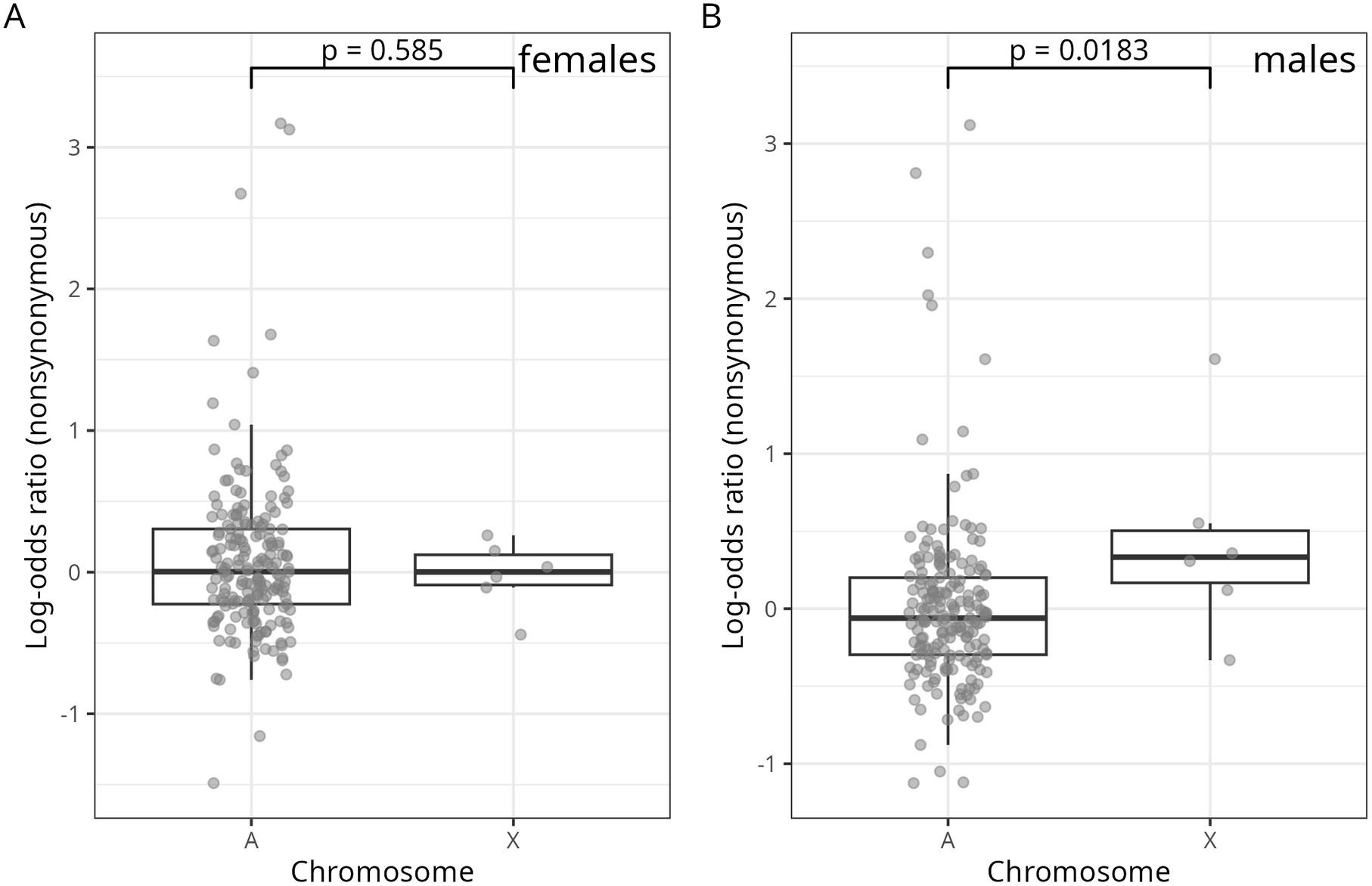
Non-synonymous mutations in OGs for males and females in autosomes and the X chromosome. Boxplot of the scaled mutation rate (log-odds ratio of the number of non-synonymous mutations, see Methods) of OGs located in any autosome (A) or in the X chromosome (X). Data is shown separately for female (A) and for male (B) cancer samples.

### Potential methodological and biological biases

To further control for methodological biases, we studied the evolutionary rates of census genes for samples that have been analysed exclusively using Whole-Genome Sequencing (WGS) and Whole-Exome Sequencing (WES), ignoring those independent studies incorporated into COSMIC. In males, the evolutionary rate of TSGs based on nonsense mutations was higher in genes in the X chromosome than in autosomes in both WGS and WES screens, while in females there was no such difference (Fig 4A and 4B). For OGs, the rate was higher in the male X in WES screens but not in WGS screens, perhaps because of the reduced number of samples.

**Fig 4.**
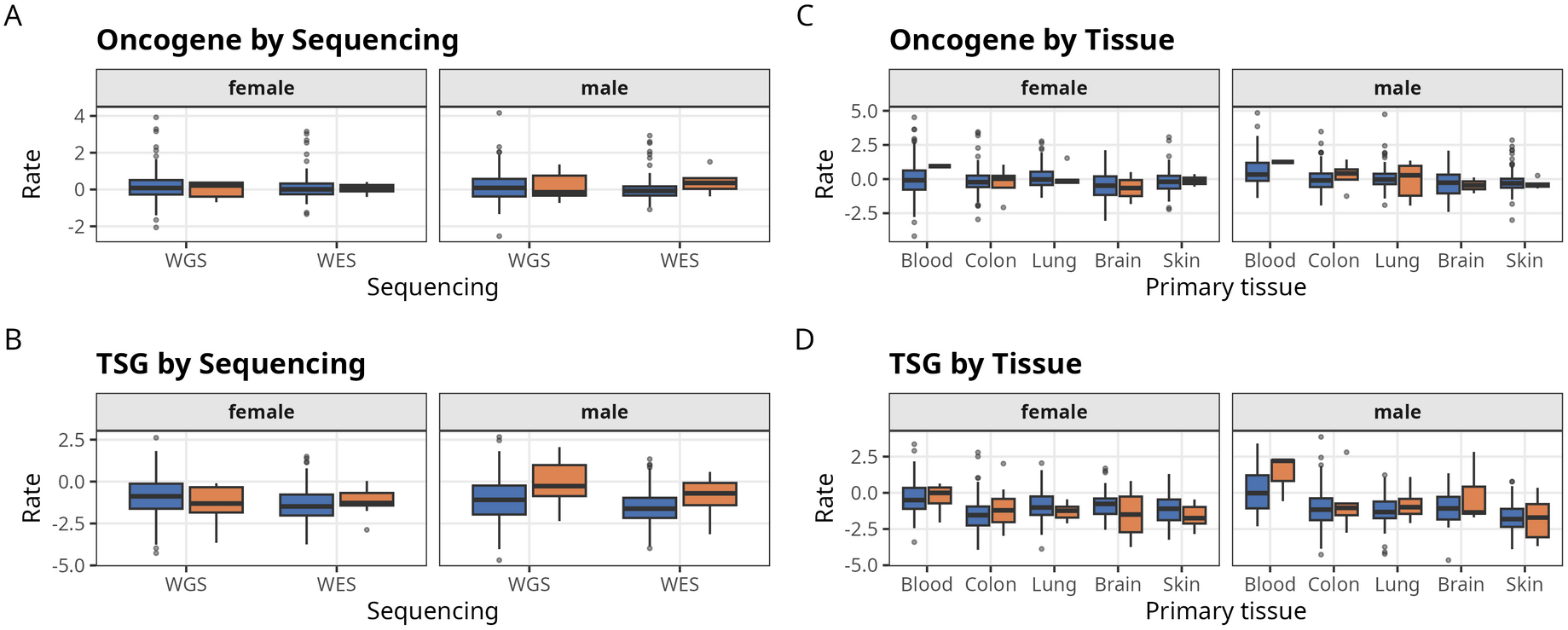
Mutations in cancer census genes pooled by genome sequencing strategy and primary tissue. Boxplots of the scaled mutation rates (see legends for Figures 2 and 3) for autosomes (blue boxes) or X chromosomes (orange boxes). The left panels show the rates for genomic screens generated by Whole-Genome Sequencing (WGS) and Whole-Exome Sequencing (WES) for OGs (A) and TSGs (B) in male and female samples. The right panels show the rates for genomic screens for five sets grouped by primary tissue: blood, colon, lung, brain and skin, for OGs (C) and TSGs (D) for male and female samples.

When we grouped the samples by primary tissue, the rate in the male X was higher than in autosomes for blood, colon and lung, but not for brain and skin (Fig 4C and 4D). This heterogeneity could reflect differences in proliferation, in the number of samples, or in the small number of X-linked genes in each group.

This raises some questions about different evolutionary rates across cancer samples depending on its biological background, possibly proliferation rates. In this respect, the model makes another prediction: for low proliferation tumours, the pattern for OGs will be reversed, and genes in the autosomes will evolve faster than genes in the X chromosome. To specifically test that, we restricted the analysis of OGs to samples from low proliferative tumours (see Methods for details). Like in the previous analysis, there is no noticeable difference in the evolutionary rates between chromosomes in female samples (Fig 5A, p = 0.216). On the other hand, and in agreement with the theoretical model, OGs in autosomes evolve faster than genes in the X although in male samples, given the limited sample size, the difference did not quite meet the significance threshold (Fig 5B, p = 0.176).

**Fig 5.**
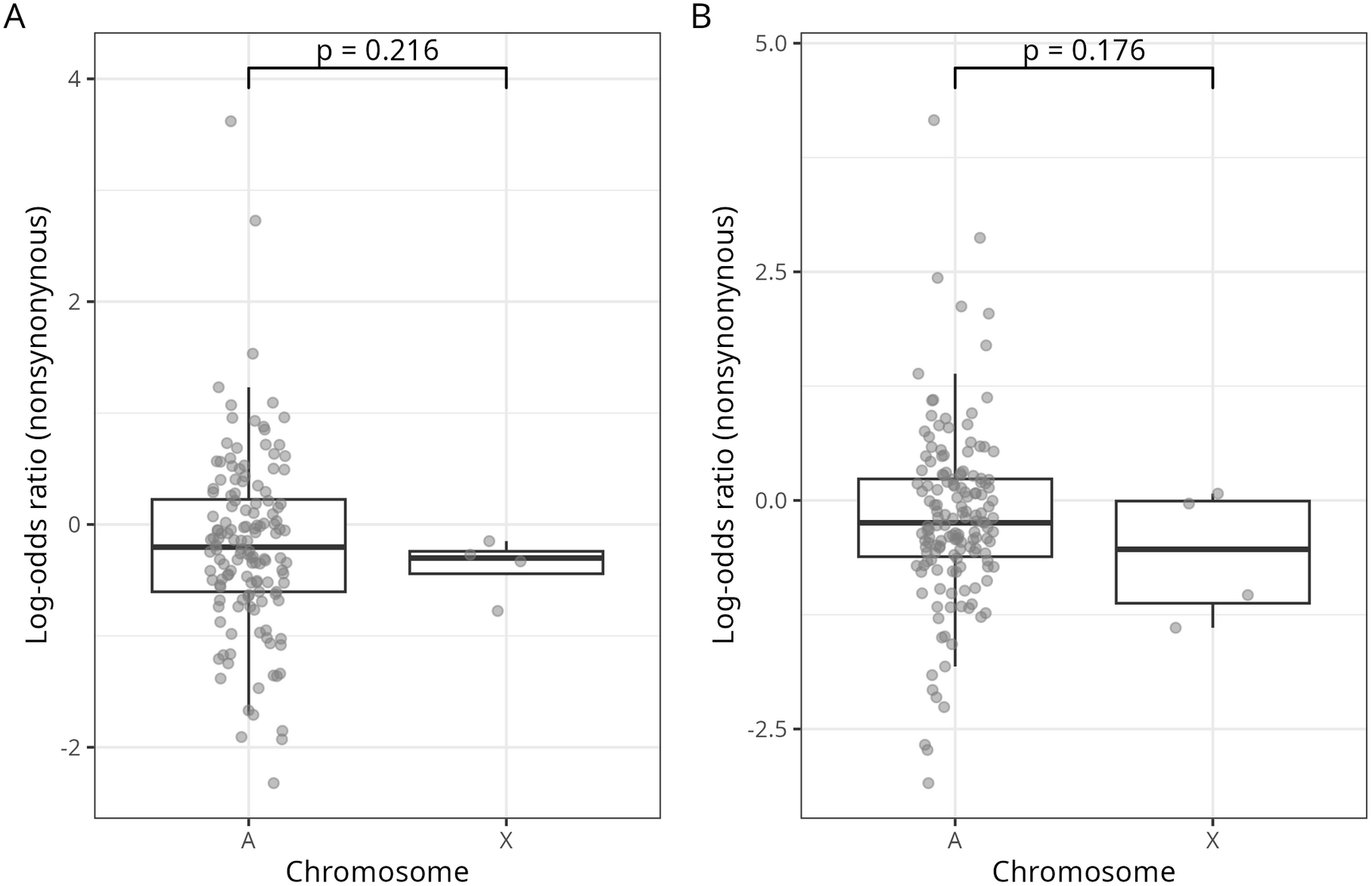
Non-synonymous mutations in low-proliferative cancer samples in OGs located in autosomes and the X chromosome. Boxplot of the scaled mutation rate (log-odds ratio of the number of non-synonymous mutations, see Methods) of OGs located in any autosome (A) or in the X chromosome (X) of low-proliferative cancer samples. Data is shown separately for female (A) and for male (B) cancer samples.

### Tumour-suppressor gene losses

In this work we have been considering loss-of-function mutations as driver mutations in TSGs, either nonsense mutations or frameshift mutations. In addition, we considered complete deletion of the TSG as another form of loss-of-function. In this context, we will also expect TSG losses to be more likely in the X chromosome than in autosomes in male samples, but not in female samples. Indeed, the number of losses in autosomes and X chromosomes in female samples is comparable (Fig 6A; p=0.548), while in male samples the number of identified TSG losses in the X chromosome is significantly higher than in autosomes (Fig 6B; p=0.001). The median number of TSG losses in X-chromosomes compared to autosomes in male samples is notable: 34%. This result complement and strengthen our observation that TSGs evolve faster in X-chromosomes compared to autosomes when accounting for various loss-of-function mutations.

**Fig 6.**
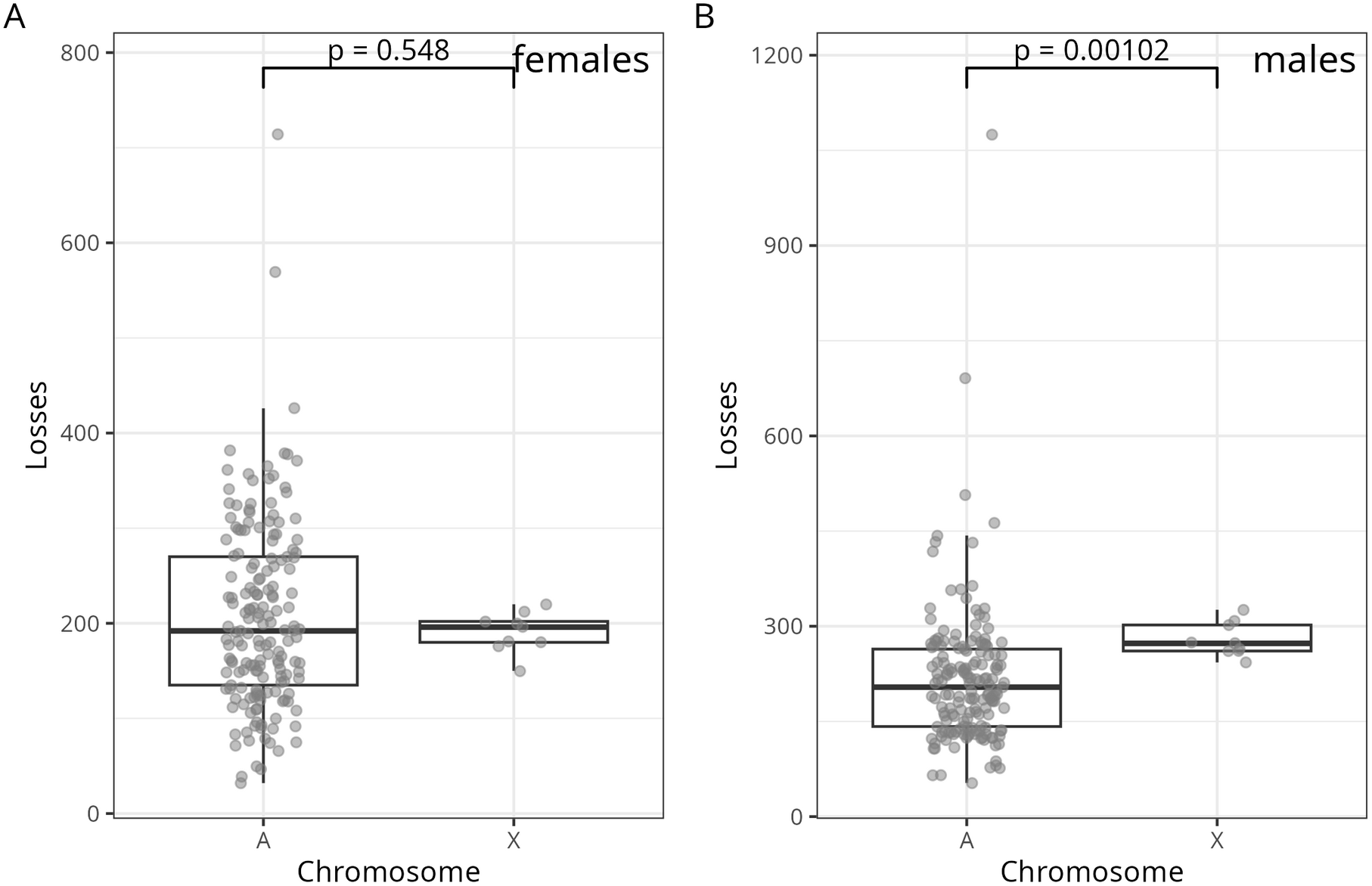
Gene losses in TSGs for males and females in autosomes and the X chromosome. Boxplot of the of the number of reported gene losses of TSGs located in any autosome (A in the x-axis) or in the X chromosome (X in the x-axis) for female (A) and for male (B) cancer samples.

### Evaluation of background neutral mutation rates

The evolutionary rates computed in this work as LOR are scaled by the number of observed and expected synonymous mutations. On the one hand, that indicates that differences in evolutionary rates are relative to the background mutation rate for each chromosome. On the other hand, differences in the mutation rate between the X chromosome and autosomes may explain some of these differences. To evaluate whether the neutral mutation rate was significantly different between chromosomes and the sex of the sample, we computed the fraction of synonymous mutations per synonymous sites and contrasted it with sex and chromosome for both TSG and OG with a linear model (regression) accounting for an interaction term (Table 2).

**Table 2.** ANOVA table (regression) of synonymous mutations for sex and chromosome.

| Type | Term | df | SS | MS | F | p |
| --- | --- | --- | --- | --- | --- | --- |
| TSG | Sex | 1 | 7.62 | 7.62 | 11.57 | 0.0007 |
|  | Chromosome | 1 | 0.50 | 0.50 | 0.76 | 0.3854 |
|  | Interaction | 1 | 1.77 | 1.77 | 2.69 | 0.1022 |
|  | Residuals | 350 | 230.59 | 0.66 |  |  |
| OG | Sex | 1 | 6.43 | 6.43 | 9.31 | 0.0024 |
|  | Chromosome | 1 | 2.07 | 2.07 | 3.00 | 0.0841 |
|  | Interaction | 1 | 3.54 | 3.54 | 5.13 | 0.0241 |
|  | Residuals | 378 | 260.96 | 0.69 |  |  |

TSG: Tumour-Suppressor Gene; OG: Oncogene; df: degrees of freedom; SS: Sum of Squares; MS: Mean Square; F: F sample statistic; p: p-value.

To build the regression model, we considered the natural logarithm of the ratio number of synonymous mutations over the number of possible synonymous mutations, as this is normally distributed (Shapiro test; p = 0.5537). For both TSGs and OGs the sex of the sample was strongly statistically associated to the synonymous mutation rate (Table 2). Specifically, male samples had higher synonymous mutation rates than female samples. However, the effect across chromosomes is not significant, supporting that our observed differences in evolutionary rates between X chromosomes and autosomes for a specific sex are not due to differences in the neutral synonymous mutation rate. We found, however, a significant interaction term for OGs, indicating that the joint effect of sex and chromosome may have an impact in the estimation of OG evolutionary rates (Table 2).

### The impact of X-chromosome Inactivation

It would be of interest to evaluate whether genes silenced by X chromosome inactivation behave as haploids in the model. However, there are two important issues. First, if we select genes that are described as X silenced or not silenced, among our TSG we only identify 4 and 5 respectively, limiting any statistical analysis. Second, gene-specific X chromosome inactivation may be very different in cancer cells compared to healthy somatic tissues, and any outcome should be taken with precaution. In any case, the median evolutionary rate (LOR) of documented X inactivated TSGs is -1.02, higher than that evolutionary rate of non-inactivated X genes and autosomes, being -1.22 and -1.50 respectively.

### Arrival times

A last prediction we made from the model is that the time of fixation of advantageous mutations is smaller in OG than in TSG except in male X chromosomes (haploids). To evaluate this we considered samples for which only mutations at either TSG or OG have been reported, and counted how many of these samples are per chromosome, sex and either TSG or OG. A shorter fixation time should be, therefore, reflected in a higher number of mutations in a set of genes compared to a gene for which a longer fixation time is expected. As expected from the model predictions, we observed a significantly higher number of mutations (shorter fixation time) in OG with respect to those in TSG for all samples except for male X chromosome genes (Table 3). That is, arrival times for (cancer) beneficial mutations in TSG and OG are comparable for X-linked genes, as predicted by the model.

**Table 3.** Expected and observed mutations found in OG and TSG.

| Chromosome | Sex | O(TSG) | E(TSG) | O(OG) | E(OG) | P<sup>1</sup> |
| --- | --- | --- | --- | --- | --- | --- |
| A | female | 370 | 1086.6 | 2968 | 2251.4 | <0.001 |
| X | female | 38 | 47.5 | 33 | 23.5 | 0.019 |
| A | male | 423 | 1156.6 | 3130 | 2396.4 | <0.001 |
| X | male | 44 | 53.6 | 36 | 26.4 | 1.000 |

<sup>1</sup>G-test (Likelihood Ratio test) associated p-values.

In summary, we showed that mathematical models of somatic evolution predict that X-linked inactivation of TSGs will evolve faster than those in the autosomes, and the available empirical data strongly support this prediction.

## DISCUSSION

Our results demonstrate that the chromosomal context and dominance relationships of mutations influence the somatic evolution of cancer genes. By combining population genetics theory with large-scale cancer genomic data, we show that the hemizygous state of the male X chromosome accelerates the accumulation of mutations in tumour-suppressor genes (TSGs), consistent with predictions derived from the Knudson’s two-hit hypothesis. In this framework, TSGs typically require inactivation of both alleles to drive tumorigenesis [6]. Our results provide support for this prediction and extend the scope of the two-hit hypothesis by incorporating chromosomal context and ploidy into models of somatic cancer evolution.

Although the model used treat cancer cells as haploid or diploid depending on chromosomal location, real tumours are often characterized by pervasive genomic instability, including whole-chromosome gains, losses, and local copy-number alterations [4]. These alterations may shift the effective ploidy of cancer genes, modulating their evolutionary dynamics. For example, loss of heterozygosity (LOH) is a well-established mechanism by which diploid TSGs effectively become hemizygous, facilitating their inactivation with a single additional mutation [6,50]. Conversely, chromosomal gains can buffer the effect of recessive mutations, delaying the fixation of loss-of-function alleles. Notably, recurrent aneuploidies are observed across multiple cancer types, and these may accelerate or decelerate the dynamics predicted by our models depending on whether they involve dosage-sensitive oncogenes (OGs) or TSGs. Indeed, recurrent copy-number alterations are particularly informative to classify cancer types [49]. However, in our genome-wide analysis of mutations, only about 5% of these alterations were in genes with a reported copy-number alteration in the same study and were excluded. Nevertheless ,incorporating variable ploidy and chromosomal instability into mathematical models therefore represents an important extension of the framework presented here.

The observation that TSGs and OGs exhibit different evolutionary dynamics could potentially be used to identify novel cancer genes, particularly TSGs, based on the differences in their mutation profiles between the X chromosome and autosomes. Also, the systematic analysis of somatic mutations in cancer genes clearly identifies that TSGs are mostly driven by nonsense mutations and OGs by missense mutations [51] in agreement with our observation (Fig 1). Indeed, the somatic evolutionary features of cancer genes have been used to classify them as either TSGs or OGs [52]. However, TSGs and OGs also have different germline evolution histories. For instance, TSGs on the X chromosome are, on average, younger than those on autosomes, suggesting that selection against TSGs on the X chromosome has driven a movement of genes out of the X [16]. This type of selection-driven movement of genes across chromosomes has been extensively studied, such as in the demasculinization of the X chromosome [53]. In summary, the distinct functional roles of TSGs and OGs in cancer are mirrored in their differing evolutionary dynamics in both germline and somatic contexts.

Our results from the analysis of genomics screens also indicated that X-chromosome inactivation is critical in cancer development, as it affects the expression of TSGs in females. Indeed, TSGs that escape X inactivation and therefore require two-hits to reach cancer status are associated with a reduction of cancer incidence in females compared to males in certain cancers [15]. Our analysis also suggest that the evolutionary rate of inactivated TSG in females is higher than that of non-inactivated (escapees) TSG in X and those in autosomes. However, the data is limited, and we could not perform adequate statistical evaluations. This is also a reflection of the complex landscape of X-chromosome inactivation in cancer as it is missing in some female cancers [54] and even some male cancers can express XIST and induce X-chromosome inactivation [55]. We additionally tested the model against low proliferative cancers (Fig 5) and, although the findings were consistent with the predictions, the statistical signal was weak. Alternative measures of cancer proliferation for larger samples may be investigated in future work.

A further limitation of our work is considering dominance solely in terms of recessive versus dominant mutations. In reality, there are cases where heterozygosity itself provides a selective advantage, a phenomenon known as overdominance or heterozygote advantage. For example, partial loss of function in DNA repair genes may increase mutation rates, fostering genetic diversity and clonal expansion, while complete loss can be deleterious to tumour cell viability [5,52]. Mathematically, overdominance is modelled by allowing values of *h* greater than 1. Thus, according to the model, overdominance will result in a systematic higher evolutionary rate of both OG and TSG in diploidy (autosomes or X in females). The available data on overdominance mutation is limited, so we could not evaluate this prediction, but we can anticipate that, if the prediction of the models are correct, future results may confirm that over dominant mutations accumulate more slowly in male X chromosomes compared to chromosomes in somatic diploidy.

Population genetics theory has been applied to somatic evolution before. The neutral theory of molecular evolution has been used to describe the dynamics of mutations in tumours [46,56], and the ratio of nonsynonymous to synonymous substitutions (dN/dS), developed for organismal evolution, has been adapted to detect selection in cancer genomes [51]. These ideas have been developed further to estimate the selective effects of individual somatic variants [46,57] and the contribution of different mutational processes to the origin of cancers [58]. Selection in tumours rarely results in hard fixation, so the concept of soft sweeps [59,60] is also relevant to the interpretation of fixation as a proxy for clonal dominance in our work. To our knowledge, however, these approaches do not model ploidy and dominance as determinants of the rate of adaptation, which is the focus of the Crow-Kimura and Orr-Otto models that we use. Our contribution is therefore not the use of population genetics in cancer, but the application of models that explicitly account for ploidy to the comparison of the X chromosome and autosomes, with separate predictions for TSGs and OGs. The statistic we use is related to dN/dS, and tools such as cancereffectsizeR [57] could provide a more powerful, context-specific estimate of selection in future work.

Taken together, our findings highlight the importance of considering chromosomal context in cancer evolution. The faster evolution of X-linked TSGs in males provides a mechanistic explanation for observed sex differences in cancer incidence and progression [7,15,61]. Moreover, our study further shows that population genetics models originally developed for organismal evolution can be adapted to somatic evolution, offering an alternative framework for understanding mutation dynamics in cancer. By extending these models to account for variable ploidy and chromosomal instability and eventually including context-dependent genetic interactions [62], we may gain deeper insights into the evolutionary logic of tumour development.

## MATERIALS AND METHODS

We retrieved information on cancer genes from the COSMIC database, version 104 (Forbes et al. 2015). We analyzed 397 census genes from Tiers 1 and 2 of COSMIC: 207 OGs and 190 TSGs, after discarding genes classified under both categories (except for the analysis in Table 1). We identified mutations described in multiple genome screens cataloged in COSMIC, tracking the individual studies and the sex of the donor. The COSMIC genomic screen collection used here comprised 82 datasets extracted from curated databases: 75 screens from database ICGC including 24 screen from TCGA and an additional 7 from PCGP. In addition, it includes 918 whole genome or systematic screens from the literature [63].

We filtered out mutations occurring in genes in which in the same sample it was reported to undergo a copy number variation, based on the CNA (copy number alterations) information form COSMIC. To filter out any gene whose methylation status may bias any measurement, we also identify the promoter region of our census genes, -2000 to 200 around the annotated transcription start site in ENSEMBL [64] and discard associated genes with a reported differential methylation in their promoters according to the COSMIC Differential Methylation information.

The log-odds ratio (LOR) was computed as:

where N_changing_ refers to either STOP gain codon (nonsense) mutations, nonsynonymous (missense) mutations or frameshift mutations, depending on the specific analysis (see main text), and in accordance N_possible_ refers to the number of possible nonsense of missense mutations. N_syn_ and N_synSites_ refer to the number of synonymous mutations and the number of synonymous sites respectively. The number of synonymous and nonsynonymous sites was computed using the Nei-Gojobori (NG86) approach [65]. The rate of synonymous somatic mutations use in Table 2 was computed as the natural logarithm of the number of synonymous mutations over the number of possible synonymous substitutions.

The primary tissues in Figure 4 corresponded to the following primary tissue samples in the COSMIC sample annotation file: ‘haematopoietic_and_lymphoid_tissue’ (blood); ‘large_intestine’ (colon); ‘lung’ (lung); ‘central_nervous_system’ (brain) and ‘skin’ (skin).

We identified studies from TCGA as low proliferative tumour samples as tumours those with a median proliferation score of under 0 according to Thorsson et al. [42], excluding only female or males tissues (prostate and breast, specifically). The TCGA studies identified as low-proliferative tumours were: KICH, PCPG, KIRP, THCA, LGG, UVM, KIRC, ACC, LIHC, PAAD, PRDA (excluded as it is prostate –male-tissue), CHOL, MESO and LUAD. X-chromosome inactivated (XCI) genes and escapees were retrieved from the compilation by Slavney et al. [43]. A full list of the census genes identified as XCI genes and escapees are listed in Supplementary Table 2.

All statistical analysis and graph plotting was done with R 4.4.0 [68] and tidyverse 2.0.0 [69]. All the code, with annotations, is available on GitHub at https://github.com/antoniomarco/CancerSomaticEvolutionX

## REFERENCES

1. Weinberg RA, Weinberg RA. The Biology of Cancer. 1st edition. New York: Garland Science; 2006.

2. Ruddon RW. Cancer Biology. Oxford University Press, USA; 2007.

3. Vogelstein B, Kinzler KW. The Genetic Basis of Human Cancer. 2nd edition. New York: McGraw-Hill Professional; 2002.

4. Greaves M, Maley CC. Clonal evolution in cancer. Nature. 2012;481: 306–313. doi:10.1038/nature10762

5. Martincorena I, Campbell PJ. Somatic mutation in cancer and normal cells. Science. 2015;349: 1483–1489. doi:10.1126/science.aab4082

6. Knudson AG. Mutation and cancer: statistical study of retinoblastoma. Proc Natl Acad Sci USA. 1971;68: 820–823.

7. Claus EB, Cannataro VL, Gaffney SG, Townsend JP. Environmental and sex-specific molecular signatures of glioma causation. Neuro Oncol. 2022;24: 29–36. doi:10.1093/neuonc/noab103

8. Rubin JB, Lagas JS, Broestl L, Sponagel J, Rockwell N, Rhee G, et al. Sex differences in cancer mechanisms. Biol Sex Differ. 2020;11: 17. doi:10.1186/s13293-020-00291-x

9. Rubin JB. The spectrum of sex differences in cancer. Trends Cancer. 2022;8: 303–315. doi:10.1016/j.trecan.2022.01.013

10. Lopes-Ramos CM, Quackenbush J, DeMeo DL. Genome-Wide Sex and Gender Differences in Cancer. Front Oncol. 2020;10: 597788. doi:10.3389/fonc.2020.597788

11. Kim H-I, Lim H, Moon A. Sex Differences in Cancer: Epidemiology, Genetics and Therapy. Biomol Ther (Seoul). 2018;26: 335–342. doi:10.4062/biomolther.2018.103

12. Haupt S, Caramia F, Klein SL, Rubin JB, Haupt Y. Sex disparities matter in cancer development and therapy. Nat Rev Cancer. 2021;21: 393–407. doi:10.1038/s41568-021-00348-y

13. Li CH, Prokopec SD, Sun RX, Yousif F, Schmitz N, PCAWG Tumour Subtypes and Clinical Translation, et al. Sex differences in oncogenic mutational processes. Nat Commun. 2020;11: 4330. doi:10.1038/s41467-020-17359-2

14. Beroukhim R, Mermel CH, Porter D, Wei G, Raychaudhuri S, Donovan J, et al. The landscape of somatic copy-number alteration across human cancers. Nature. 2010;463: 899–905. doi:10.1038/nature08822

15. Dunford A, Weinstock DM, Savova V, Schumacher SE, Cleary JP, Yoda A, et al. Tumor-suppressor genes that escape from X-inactivation contribute to cancer sex bias. Nat Genet. 2017;49: 10–16. doi:10.1038/ng.3726

16. Wang X, Hu W, Li X, Huang D, Li Q, Chan H, et al. Single-Hit Inactivation Drove Tumor Suppressor Genes Out of the X Chromosome during Evolution. Cancer Research. 2022;82: 1482–1491. doi:10.1158/0008-5472.CAN-21-3458

17. Ryland GL, Doyle MA, Goode D, Boyle SE, Choong DYH, Rowley SM, et al. Loss of heterozygosity: what is it good for? BMC Med Genomics. 2015;8: 45. doi:10.1186/s12920-015-0123-z

18. Dutta A, Schacherer J. The dynamics of loss of heterozygosity events in genomes. EMBO Rep. 2025;26: 602–612. doi:10.1038/s44319-024-00353-w

19. Frank SA. Dynamics of Cancer: Incidence, Inheritance, and Evolution. Princeton University Press; 2018.

20. Frank SA. Somatic evolutionary genomics: Mutations during development cause highly variable genetic mosaicism with risk of cancer and neurodegeneration. Proceedings of the National Academy of Sciences. 2010;107: 1725–1730. doi:10.1073/pnas.0909343106

21. Nunney L. Lineage selection and the evolution of multistage carcinogenesis. Proc Biol Sci. 1999;266: 493–498.

22. Frank SA, Nowak MA. Problems of somatic mutation and cancer. Bioessays. 2004;26: 291–299. doi:10.1002/bies.20000

23. Michor F, Frank SA, May RM, Iwasa Y, Nowak MA. Somatic selection for and against cancer. J Theor Biol. 2003;225: 377–382. doi:10.1016/s0022-5193(03)00267-4

24. Alfaro-Murillo JA, Townsend JP. Pairwise and higher-order epistatic effects among somatic cancer mutations across oncogenesis. Mathematical Biosciences. 2023;366: 109091. doi:10.1016/j.mbs.2023.109091

25. Cairns J. Mutation and cancer: the antecedents to our studies of adaptive mutation. Genetics. 1998;148: 1433–1440. doi:10.1093/genetics/148.4.1433

26. Crow JF, Kimura M. An Introduction to Population Genetics Theory. Harper & Row; 1970.

27. Hartl DL, Clark AG. Principles of Population Genetics. Sinauer Associates; 2007.

28. Charlesworth B, Coyne JA, Barton NH. The Relative Rates of Evolution of Sex Chromosomes and Autosomes. The American Naturalist. 1987;130: 113–146. doi:10.2307/2461884

29. Avila V, de Procé SM, Campos JL, Borthwick H, Charlesworth B, Betancourt AJ. Faster-X effects in two Drosophila lineages. Genome Biol Evol. 2014. doi:10.1093/gbe/evu229

30. Fraïsse C, Puixeu Sala G, Vicoso B. Pleiotropy Modulates the Efficacy of Selection in Drosophila melanogaster. Mol Biol Evol. 2019;36: 500–515. doi:10.1093/molbev/msy246

31. Crow JF, Kimura M. Evolution in Sexual and Asexual Populations. The American Naturalist. 1965;99: 439–450. doi:10.1086/282389

32. Fisher RA. The genetical theory of natural selection. Oxford Clarendon Press; 1930. Available: http://archive.org/details/geneticaltheoryo00fishuoft

33. Muller HJ. Some Genetic Aspects of Sex. The American Naturalist. 1932;66: 118–138. doi:10.1086/280418

34. Orr HA, Otto SP. Does diploidy increase the rate of adaptation? Genetics. 1994;136: 1475–1480. doi:10.1093/genetics/136.4.1475

35. Haldane JBS. A Mathematical Theory of Natural and Artificial Selection. Transactions of the Cambridge Philosophical Society. 1924;23: 19–41.

36. Knudson AG. Mutation and Cancer: Statistical Study of Retinoblastoma. Proc Natl Acad Sci U S A. 1971;68: 820–823.

37. Milholland B, Dong X, Zhang L, Hao X, Suh Y, Vijg J. Differences between germline and somatic mutation rates in humans and mice. Nat Commun. 2017;8: 15183. doi:10.1038/ncomms15183

38. Ju YS, Martincorena I, Gerstung M, Petljak M, Alexandrov LB, Rahbari R, et al. Somatic mutations reveal asymmetric cellular dynamics in the early human embryo. Nature. 2017;543: 714–718. doi:10.1038/nature21703

39. Lynch M. Rate, molecular spectrum, and consequences of human mutation. Proceedings of the National Academy of Sciences. 2010;107: 961–968. doi:10.1073/pnas.0912629107

40. Weinstein GD, McCullough JL, Ross P. Cell proliferation in normal epidermis. J Invest Dermatol. 1984;82: 623–628. doi:10.1111/1523-1747.ep12261462

41. Yu C-Y, Lin C-H, Yang Y-H. Human body surface area database and estimation formula. Burns. 2010;36: 616–629. doi:10.1016/j.burns.2009.05.013

42. Wright JJ, Saunders DC, Dai C, Poffenberger G, Cairns B, Serreze DV, et al. Decreased pancreatic acinar cell number in type 1 diabetes. Diabetologia. 2020;63: 1418–1423. doi:10.1007/s00125-020-05155-y

43. Grimont A, Leach SD, Chandwani R. Uncertain Beginnings: Acinar and Ductal Cell Plasticity in the Development of Pancreatic Cancer. Cell Mol Gastroenterol Hepatol. 2022;13: 369–382. doi:10.1016/j.jcmgh.2021.07.014

44. Jiang Z, White RA, Wang TC. Adult Pancreatic Acinar Progenitor-like Populations in Regeneration and Cancer. Trends Mol Med. 2020;26: 758–767. doi:10.1016/j.molmed.2020.04.003

45. Wollny D, Zhao S, Everlien I, Lun X, Brunken J, Brüne D, et al. Single-Cell Analysis Uncovers Clonal Acinar Cell Heterogeneity in the Adult Pancreas. Developmental Cell. 2016;39: 289–301. doi:10.1016/j.devcel.2016.10.002

46. Cannataro VL, Townsend JP. Neutral Theory and the Somatic Evolution of Cancer. Mol Biol Evol. 2018;35: 1308–1315. doi:10.1093/molbev/msy079

47. Gedye C, Sirskyj D, Lobo NC, Meens J, Hyatt E, Robinette M, et al. Cancer stem cells are underestimated by standard experimental methods in clear cell renal cell carcinoma. Sci Rep. 2016;6: 25220. doi:10.1038/srep25220

48. Grimmett G, Stirzaker D. Probability and Random Processes: Fourth Edition. Oxford: Oxford University Press; 2020.

49. Iwasa Y, Michor F, Komarova NL, Nowak MA. Population genetics of tumor suppressor genes. Journal of Theoretical Biology. 2005;233: 15–23. doi:10.1016/j.jtbi.2004.09.001

50. Ciriello G, Miller ML, Aksoy BA, Senbabaoglu Y, Schultz N, Sander C. Emerging landscape of oncogenic signatures across human cancers. Nat Genet. 2013;45: 1127–1133. doi:10.1038/ng.2762

51. Martincorena I, Raine KM, Gerstung M, Dawson KJ, Haase K, Van Loo P, et al. Universal Patterns of Selection in Cancer and Somatic Tissues. Cell. 2017;171: 1029–1041.e21. doi:10.1016/j.cell.2017.09.042

52. Chandrashekar P, Ahmadinejad N, Wang J, Sekulic A, Egan JB, Asmann YW, et al. Somatic selection distinguishes oncogenes and tumor suppressor genes. Bioinformatics. 2020;36: 1712–1717. doi:10.1093/bioinformatics/btz851

53. Sturgill D, Zhang Y, Parisi M, Oliver B. Demasculinization of X chromosomes in the Drosophila genus. Nature. 2007;450: 238–241. doi:10.1038/nature06330

54. Cáceres A, Pérez-Jurado LA, Alegret-García A, Dwaraka VB, Smith R, González JR. Defective X-chromosome inactivation and cancer risk in women. Commun Biol. 2025;8: 289. doi:10.1038/s42003-025-07691-y

55. Sadagopan A, Nasim IT, Li J, Achom M, Zhang C-Z, Viswanathan SR. Somatic XIST activation and features of X chromosome inactivation in male human cancers. cels. 2022;13: 932–944.e5. doi:10.1016/j.cels.2022.10.002

56. Niida A, Iwasaki WM, Innan H. Neutral Theory in Cancer Cell Population Genetics. Mol Biol Evol. 2018;35: 1316–1321. doi:10.1093/molbev/msy091

57. Mandell JD, Cannataro VL, Townsend JP. Estimation of Neutral Mutation Rates and Quantification of Somatic Variant Selection Using cancereffectsizeR. Cancer Res. 2023;83: 500–505. doi:10.1158/0008-5472.CAN-22-1508

58. Cannataro VL, Mandell JD, Townsend JP. Attribution of Cancer Origins to Endogenous, Exogenous, and Preventable Mutational Processes. Mol Biol Evol. 2022;39: msac084. doi:10.1093/molbev/msac084

59. Pritchard JK, Pickrell JK, Coop G. The genetics of human adaptation: hard sweeps, soft sweeps, and polygenic adaptation. Curr Biol. 2010;20: R208–215. doi:10.1016/j.cub.2009.11.055

60. Messer PW, Petrov DA. Population genomics of rapid adaptation by soft selective sweeps. Trends in Ecology & Evolution. 2013;28: 659–669. doi:10.1016/j.tree.2013.08.003

61. Haupt S, Caramia F, Klein SL, Rubin JB, Haupt Y. Sex disparities matter in cancer development and therapy. Nat Rev Cancer. 2021;21: 393–407. doi:10.1038/s41568-021-00348-y

62. Chang L, Shaw K, Vazquez F, Sellers WR. Context-dependent synthetic lethality — an emerging precision therapeutic approach. Nat Rev Cancer. 2026;26: 534–554. doi:10.1038/s41568-026-00929-9

63. Forbes SA, Beare D, Gunasekaran P, Leung K, Bindal N, Boutselakis H, et al. COSMIC: exploring the world’s knowledge of somatic mutations in human cancer. Nucl Acids Res. 2014; gku1075. doi:10.1093/nar/gku1075

64. Kinsella RJ, Kähäri A, Haider S, Zamora J, Proctor G, Spudich G, et al. Ensembl BioMarts: a hub for data retrieval across taxonomic space. Database (Oxford). 2011;2011: bar030. doi:10.1093/database/bar030

65. Nei M, Gojobori T. Simple methods for estimating the numbers of synonymous and nonsynonymous nucleotide substitutions. Mol Biol Evol. 1986;3: 418–426. doi:10.1093/oxfordjournals.molbev.a040410

66. Thorsson V, Gibbs DL, Brown SD, Wolf D, Bortone DS, Ou Yang T-H, et al. The Immune Landscape of Cancer. Immunity. 2018;48: 812–830.e14. doi:10.1016/j.immuni.2018.03.023

67. Slavney A, Arbiza L, Clark AG, Keinan A. Strong Constraint on Human Genes Escaping X-Inactivation Is Modulated by their Expression Level and Breadth in Both Sexes. Mol Biol Evol. 2016;33: 384–393. doi:10.1093/molbev/msv225

68. R Development Core Team. R: A language and environment for statistical computing. 2004 [cited 1 May 2013]. Available: http://www.R-project.org

69. Wickham H, Averick M, Bryan J, Chang W, McGowan LD, François R, et al. Welcome to the Tidyverse. Journal of Open Source Software. 2019;4: 1686. doi:10.21105/joss.01686

